# Sex chromosomes shape the transcriptional landscape of the preimplantation mouse embryo

**DOI:** 10.1101/2024.11.26.625424

**Authors:** Daniel M. Snell, Wazeer Varsally, Aurélien Courtois, Sergio Menchero, Prabhakaran Munusamy, Richelle Rietdijk, Obah A. Ojarikre, Stephanie Strohbuecker, Haskan Kaya, Mahesh N. Sangrithi, James M.A. Turner

## Abstract

Sex chromosomes are emerging as key regulators of adult health and disease in males (XY) and females (XX), but their impact on embryo development is poorly understood. Using single-cell RNA sequencing (scRNA-seq) on wild type and aneuploid mouse embryos, we show that sex chromosomes significantly shape the preimplantation embryo transcriptional landscape. A hierarchy of effects are identified, distinctly mediated by the Y chromosome, the dosage of X chromosomes, X-chromosome imprinting, and by *Xist*, the non-coding RNA that initiates X-inactivation. The sex chromosomes have strong *trans* effects on autosomal gene expression throughout preimplantation development. The Y chromosome has an unexpectedly pronounced impact on the trophectoderm, the precursor of the placenta, and this property is shared with genes expressed from the inactive X chromosome. The paternal and maternal X chromosomes differentially promote preimplantation growth, and we identify multiple novel candidate X-linked imprinted genes mediating this effect. Our findings show that sex chromosomes impact the embryo from the beginning of life, long before the appearance of overt sex differences.

## Introduction

Sex chromosomes play a crucial role in a wide range of biological processes, affecting gene activity and protein production at the cellular level to an organism’s anatomy, physiology, and disease risk. This influence is evident in the many physical differences between males and females across species and in the phenotypes seen in individuals with atypical numbers of sex chromosomes^1–6^. Most research has focused on the impact of sex chromosomes later in development or in adulthood. However, morphological studies suggest that sex chromosomes also affect the early embryo. For example, mouse^7–10^, rat^11^, cow^12,13^ and human^14,15^ XY embryos are developmentally advanced relative to XX embryos. Additionally, at the post-implantation stage, embryos with a single X chromosome of maternal origin (X^m^O) are developmentally advanced relative to those with a single X of paternal origin (X^p^O). Thus, the X chromosome carries as-yet undiscovered imprinted genes that regulate developmental pace^8,9,16^. These sex chromosome effects are evident before gonad formation, and are thus entirely independent of sex hormones.

We currently lack insight into these sex chromosome effects, because transcriptional analysis of embryos with distinct sex chromosome complements has not been performed. The preimplantation period is of particular interest, since X-dosage compensation is not yet complete, thereby allowing sex chromosomes to exert a stronger influence on sex differences. X-dosage compensation is critical for embryo development, and in mice is achieved by two collaborative mechanisms. The first, X-chromosome upregulation, occurs in both males and females, and ensures that gene expression from the single active X chromosome is balanced with that from the autosomes^17–19^. The second, X-chromosome inactivation, occurs only in females, and ensures that gene expression from the two X chromosomes is balanced with the single X in males^20,21^. X-inactivation is initiated by the long non-coding RNA *Xist*^21–23^. In early mouse embryos, X-inactivation is imprinted, with the paternal X chromosome chosen for silencing^24–26^. Imprinted X-inactivation initiates from the four-cell stage and is retained in the trophectoderm, the precursor of the placenta. However, it is reversed in the epiblast, the cell population that will give rise to the embryo proper, where random X-inactivation then occurs^25–28^. How X-dosage compensation plays out in embryos with atypical sex chromosome complements is not well understood.

Here, we use mouse models carrying distinct sex chromosome combinations to reveal the impact of the Y chromosome, X-dosage, X-imprinting and *Xist* on mouse preimplantation development.

## Results

### An scRNA-seq dataset of sex chromosome variant embryos

To determine how sex chromosomes impact embryonic gene expression, we performed scRNA-seq on embryonic day (E) 2.0, 2.5 and 3.5 embryos derived from well-established parental sex chromosome variant lines^6,29–31^. In our colony these timepoints correspond to 4-cell, morula and blastocyst stage embryos, respectively. From a larger cohort of embryos, we extracted genotypes carrying two X chromosomes, one X and one Y chromosome, one X chromosome of maternal origin, and one X chromosome of paternal origin (hereafter denoted as X^m^X^p^, X^m^Y, X^m^O and X^p^O; Extended Data Fig. 1a,b; Methods). Comparing these genotypes allowed us to identify X^m^X^p^ - X^m^Y differences, Y-dosage and X-dosage effects, and X-imprinting effects (Fig. 1a). We ensured that the proportions of each genotype were consistent at each gestational age (Extended Data Fig. 1c). 1595 single cells (generated from a total of 191 embryos), with an average of 40,358 expressed genes, were retained after quality control. Uniform manifold approximation and projection (UMAP) revealed that samples separated according to gestational age, and that the expected trophectoderm and inner cell mass populations were apparent at E3.5 (Fig 1b; see later for exceptional genotypes).

**Figure 1.**
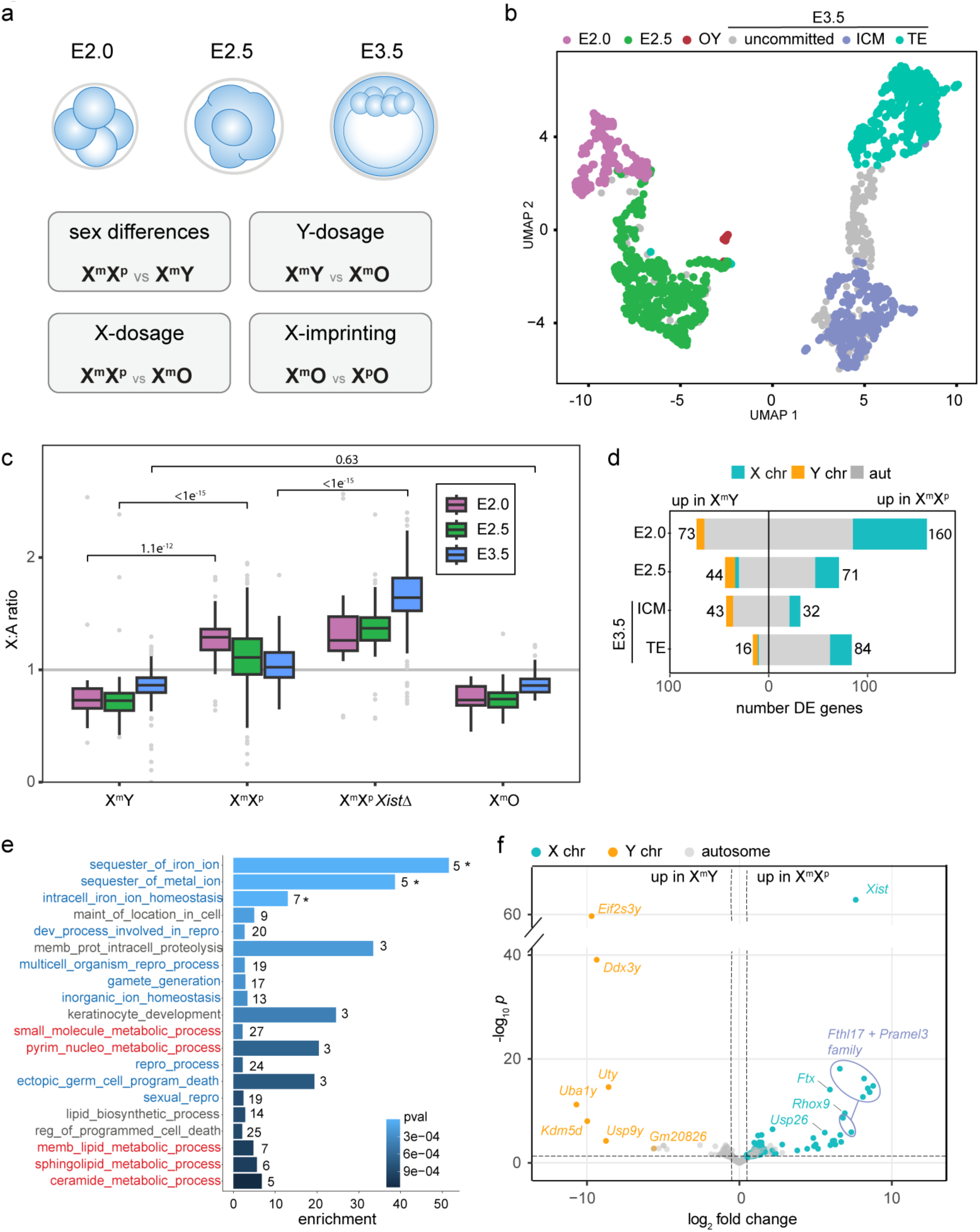
X^m^X^p^ - X^m^Y differences in pre-implantation mouse embryos. **a**. Cartoon depicting embryo stages and representing which genotypes were used to assess X^m^X^p^ - X^m^Y differences, Y-dosage, X-dosage and X imprinting effects. **b**. UMAP annotated by clusters relating to collection timepoint (E2.0, E2.5), cell lineage identified by marker analysis (E3.5 broken into inner cell mass (ICM), trophectoderm (TE), and uncommitted), or genotype (OY). **c**. Comparison of X-to-autosome expression by X:A ratios; *P* values derived from Wilcoxon test. **d**. Bar plot of DE genes between X^m^X^p^ and X^m^Y embryos. **e**. Functional enrichment analysis of X^m^X^p^ vs X^m^Y E2.0 DE genes, with terms comprising genes with putative reproductive functions (including the iron-binding *Fthl17* genes) highlighted in blue, and metabolic genes highlighted in red; * = adjusted *P* value <0.05. **f**. Volcano plot of E2.0 DE genes (note split Y axis). Horizontal dashed line represents transformed *P* value = 0.05. Vertical dashed line represents log_2_FC = 0.5.

### XX-XY differences

We examined gene expression differences between X^m^Y and X^m^X^p^ embryos. We focused initially on X-dosage compensation, which was assayed using X-to-autosome (X:A) ratio calculations (Fig. 1c). In males, X-upregulation balances X to autosomal output by increasing the X:A ratio from 0.5 to 1.0. X-upregulation initiated early in X^m^Y males, with the X:A ratio already >0.5 at E2.0 and approaching 1.0 by E3.5. A similar profile was observed in X^m^O embryos, demonstrating that X-upregulation occurs independent of the presence of a second sex chromosome. In X^m^X^p^ female embryos, X-upregulation also initiated early, with the X:A ratio >1.0 at E2.0 and E2.5. As a result, the X:A ratio was significantly higher in X^m^X^p^ than X^m^Y embryos at these timepoints. Subsequently, the X:A ratio in X^m^X^p^ embryos decreased towards 1.0. This decline was due to X-inactivation, because it did not occur in X^m^X^p^ females carrying a paternal *Xist* deletion (Fig. 1c)^21^. Our findings are consistent with reports that X-upregulation precedes X-inactivation during mouse preimplantation embryo development^18,19,32,33^.

We examined differential gene expression between X^m^Y and X^m^X^p^ embryos. The number of differentially expressed (DE) genes was highest at E2.0 and thereafter decreased (Fig. 1d). Further examination of DE genes revealed notable findings. X-genes were over-represented on a *per chromosome* basis at all timepoints (Extended Data Fig. 2a), presumably because X-dosage compensation was not yet complete. However, most DE genes were autosomal. Gene ontology analysis revealed distinct pathways enriched at successive stages of development (E2.0: Fig. 1e; E2.5 onwards Extended Data Fig. 2b; Supplementary Table 1). Among the E2.0 enriched pathways, we observed several categories related to metabolism and reproduction (Fig. 1e). We expected that Y-genes would account for the reproduction categories, because the Y chromosome is specialised for the male germline^34^. Indeed, the most strongly male-biased genes included six of the 14 genes on the mouse Y chromosome short arm; *Eif2s3y, Ddx3y, Uty, Kdm5d, Uba1y* and *Usp9y* (Fig. 1f; Extended Data Fig. 2c). These Y-genes encode proteins with dosage-sensitive housekeeping functions, and two (*Uba1y* and *Usp9y*) exhibit testis-biased expression^35–38^. Strongly female-biased genes included *Xist* and its upstream regulator *Ftx*, which are imprinted and expressed only from X^p^ ^39,40^. However, they also included members of X-linked multicopy families that were previously described as testis-biased, including the *Fthl17* iron-binding, *Rhox* homeobox and *Pramel3* transcription regulator genes^41,42^, and others (Fig. 1f; see later). We conclude that sex chromosomes act *in trans* to create sexually dimorphic autosomal expression profiles in the early embryo, and that some sex-linked genes with canonical reproductive functions are also expressed at this time.

### The impact of Y- and X-dosage

To examine how the expression of Y-genes in the early embryo influences the preimplantation transcriptome, we compared gene expression between X^m^Y and X^m^O embryos (Fig. 2a). A relatively modest effect on gene expression was observed at E2.0 and E2.5. However, there was a disproportionately high number of DE genes at E3.5 within the trophectoderm (420 genes; compared with 57 in the inner cell mass). This finding revealed an unexpected role for the Y chromosome in regulating the trophectoderm transcriptome. When we compared X^m^O to X^m^X^p^ embryos, we observed a similar trophectoderm-biased effect (444 DE genes; compared with 45 in the inner cell mass; Fig. 2b). Thus, the trophectoderm is uniquely sensitive to loss of either the Y chromosome or the paternal inactive X chromosome.

**Figure 2.**
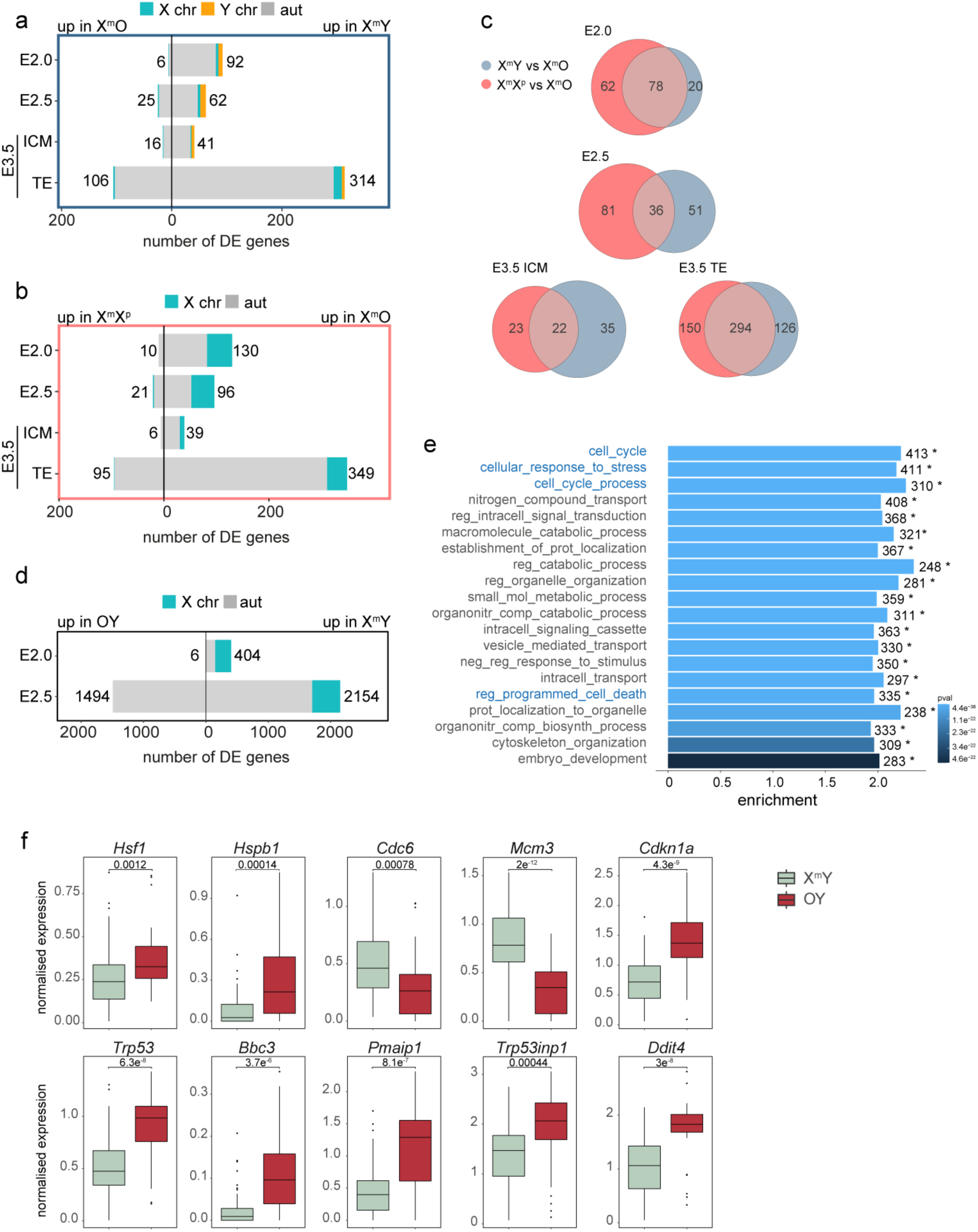
Impact of Y-dosage and X-dosage. **a**. Bar plot of DE genes between X^m^Y and X^m^O embryos. **b**. Bar plot of DE genes between X^m^X^p^ vs. X^m^O embryos. **c**. Overlap of DE genes between X^m^Y vs. X^m^O and X^m^X^p^ vs. X^m^O comparisons. **d**. Bar plot of DE genes between X^m^Y and OY embryos. **e**. Functional enrichment analysis of DE genes comparing X^m^Y and OY embryos at E2.5, with terms of specific interest highlighted in blue; * = adjusted *P* value <0.05. **f**. Normalised expression of genes highlighted in main text, comparing X^m^Y and OY embryos at E2.5; *P* values derived from Student’s t-test.

We hypothesised that the trophectoderm haploinsufficiency effect was mediated by genes shared between the Y and paternal inactive X. In support of this conclusion, there was a large overlap in DE genes between the trophectoderm cells of X^m^Y and X^m^X^p^ compared to X^m^O embryos (Fig. 2c). This effect is likely mediated by X-Y homologous gene pairs in which the X-copy escapes X-inactivation, resulting in a double dose of sex-products in both sexes (X + Y in males, X + X in females). Indeed, four of the Y-genes we found expressed in embryos: *Uty, Kdm5d, Ddx3y* and *Eif2s3y*, have X-linked homologues that escape X-inactivation^36^. Functional enrichment analysis revealed that these shared targets span many biological pathways (Extended Data Fig. 3), presumably because X^m^O mice are simultaneously haploinsufficient for multiple, dosage-sensitive X-Y homologues. The Y chromosome and paternal inactive X chromosome therefore impart overlapping effects on trophectoderm transcription.

Our sex chromosome variant crosses also generated OY embryos, which carry no X chromosome. By comparing these to X^m^Y embryos, we were able to reveal for the first time the transcriptional effect of having no X chromosome. Previous work has shown that OY embryos arrest at the 8-cell stage^43^, but whether this results directly from a lack of X-linked transcripts or indirectly from a *trans* effect of X-genes on autosomes is unclear. On the UMAP, OY embryos formed a separate cluster from other genotypes (Fig. 1b). At E2.0, most genes differentially expressed between OY and X^m^Y embryos were X-linked; however, by E2.5, we observed strong changes in autosomal gene expression (Fig. 2d). This finding further emphasised the *trans*-effect of X-genes on autosomes. Functional enrichment analysis at E2.5 revealed several pathways affected by absence of the X, with the most significant being those related to cellular stress responses, the cell cycle, and apoptosis (Fig. 2e). OY embryos exhibited upregulation of aneuploidy-associated proteotoxic response genes, such as *Hsf1* and *Hspb1*, and downregulation of replication licensing genes *Cdc6* and *Mcm3,* suggestive of replicative stress (Fig. 2f)^44^. They also exhibited upregulation of *Cdkn1a*, a key regulator of the cell cycle^45,46^, and the apoptotic factor *Trp53*, along with its downstream pro-apoptotic BCL-2 family targets *Bbc3* (*Puma*) and *Pmaip1* (*Noxa*; Fig. 2f). Additionally, genes related to autophagy and mitophagy, such as *Trp53inp1* and *Ddit4*^47^ were upregulated (Fig. 2f). Our findings suggest that embryos lacking an X exhibit signs of aneuploidy-induced stress, associated with cell cycle arrest and apoptosis.

### The impact of X-imprinting

To establish X-imprinting effects on preimplantation development, we then analysed the X^p^O model. Previous work showed that X^p^O embryos are developmentally delayed relative to X^m^O, X^m^Y and X^m^X^p^ embryos at E7.5 and E10.5 ^9,16^. We discovered that developmental delay was also evident in X^p^O embryos before implantation. Using immunostaining for markers of the inner cell mass (NANOG) and trophectoderm (CDX2), we found that X^p^O embryos had fewer cells than X^m^Y and X^m^X^p^ embryos at E3.5 (Fig. 3a). Furthermore, while E3.5, X^m^Y, X^m^X^p^ and X^m^O cells segregated into trophectoderm and inner cell mass, E3.5 X^p^O embryos exhibited transcriptional delay, with a higher proportion of cells mapping to the uncommitted cluster and a lower proportion of cells mapping to the inner cell mass cluster (Fig. 3b,c; Fig. 1b). The trophectoderm of the X^p^O embryo also exhibited developmental delay, expressing wild type levels of the early trophectoderm marker *Cdx2* but lower levels of late trophectoderm markers *Elf5* and *Eomes*^48^ (Fig. 3a,d).

**Figure 3.**
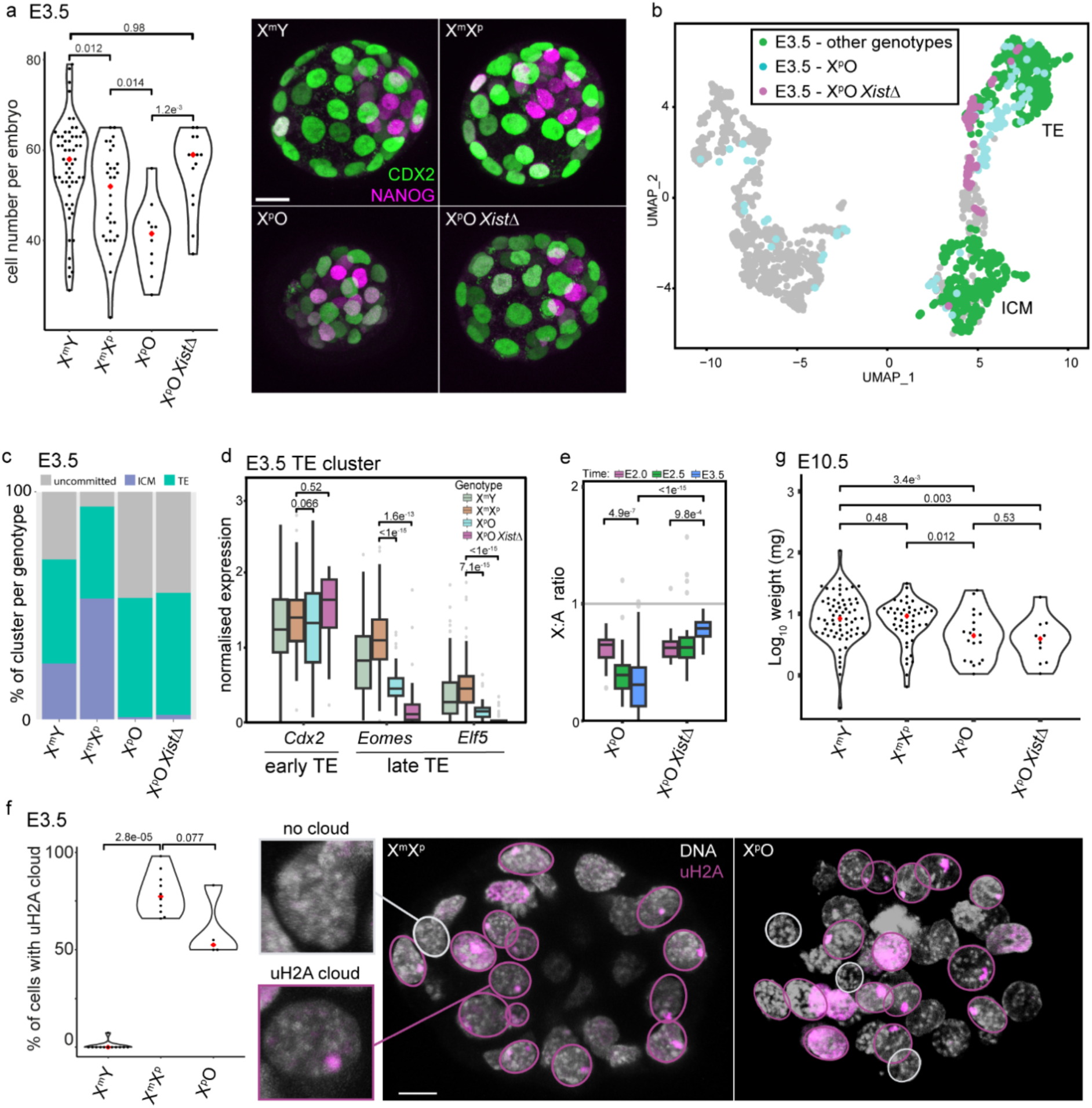
Impact of X-imprinting. **a**. Left: total cell number in E3.5 embryos (X^m^Y n=57, X^m^X^p^ n= 31, X^p^O=10, X^p^O *Xist*^Δ^ n= 14 embryos); mean value indicated by a red diamond; *p*-value generated using Student’s t-test. Right: Z-projection of confocal sections (1µm thickness) of E3.5 embryos stained for CDX2 (outside cells), and NANOG (inside cells); scale bar: 20µm. **b**. UMAP showing E3.5 X^p^O (blue) and X^p^O *Xist*^Δ^ (magenta) cells relative to all other genotypes at E3.5 (green) and other time points (grey). **c**. Proportion of cells within each cluster (uncommitted, ICM, TE) per genotype at E3.5, derived from UMAP in **b**. **d**. Normalised expression of early (*Cdx2*) and late (*Eomes*, *Elf5*) TE genes in E3.5 TE cluster cells. **e**. X-to-autosome (X:A) ratios of X^p^O and X^p^O *Xist*^Δ^ embryos; *P* values derived from Student’s t-test; one point was excluded as it is outside the represented range. **f**. Left: quantitation of inactive X-associated ubiquitylated histone H2A at lysine-119 (uH2A) staining in E3.5 embryos (X^m^Y n=12, X^m^X^p^ n=10, X^p^O n=4 embryos); mean value indicated by a red diamond. Right: Z-projection of confocal sections (1µm thickness – total thickness of 10 and 4.5µm respectively) of E3.5 embryos showing uH2A staining. Presence and absence of inactive X-associated uH2A shown by magenta and white circles, respectively. Inserts show a representative image of cells without and with a uH2A cloud. **g**. E10.5 embryo weights in mg (log_10_) by genotypes; mean value indicated by a red diamond; *P* values derived from Wilcoxon test.

Two mechanisms could contribute to this delay. One possibility is that X-expression in X^p^O embryos is globally reduced by imprinted X-inactivation, leading to a paucity of X-gene products. Indeed, X^p^O embryos exhibit imprinted *Xist* expression from the 8-cell to the blastocyst stage^49^, but whether this triggers X-inactivation of the single X is unknown. A second possibility is an *Xist*-independent effect, mediated by a growth repressor expressed only from the paternal X, and/or growth promoter expressed only from the maternal X.

To investigate the X-inactivation model, we assayed X-expression in X^p^O embryos. We found that X-inactivation was indeed initiated. This was evident from X:A ratios, which decreased during X^p^O preimplantation development (Fig. 3e). Using RNA-FISH, we confirmed that X^p^O embryos express *Xist*. Furthermore, they initiated silencing of the X-linked gene *Atp7a*, which has been shown to be inactivated by *Xist* early in female development (Extended Data Fig. 4)^50^. The single X chromosome in X^p^O embryos also accumulated ubiquitylation of histone H2A at lysine-119, an *Xist*-dependent repressive epigenetic mark indicative of X-inactivation (Fig. 3f)^51,52^.

To resolve whether the growth delay was *Xist* dependent, we generated X^p^O embryos carrying a paternal *Xist* deletion^21^. In the resulting X^p^O *Xist*^Δ^ embryos, the X:A ratio increased during embryo development (Fig. 3e). This behaviour mirrored that observed in X^m^O embryos (Fig. 1c). The paternal X is therefore competent for upregulation if X-inactivation is abolished. Notably, *Xist* deletion rescued the low blastomere count phenotype in X^p^O embryos at E3.5 (Fig. 3a). However, it did not rescue the transcriptional delay. E3.5 X^p^O *Xist*^Δ^ embryos exhibited a similar phenotype to X^p^O embryos, with a higher proportion of uncommitted cells, a lower proportion of inner cell mass cells, and lower expression of late trophectoderm markers (Fig. 3b-d). Thus, an *Xist*-independent X-imprinting effect influences development of the preimplantation embryo.

We next established whether this *Xist*-independent effect was also responsible for the growth delay originally observed in X^p^O embryos later in gestation, at E10.5. Indeed, X^p^O *Xist*^Δ^ embryos were growth delayed at E10.5, being of comparable size to X^p^O embryos and significantly smaller than their wild type counterparts (Fig. 3g). We conclude that X-parent-of-origin influences pre- and post-implantation mouse development in a manner that is independent of *Xist*.

### Identification of candidate preimplantation X-imprinted genes

Motivated by our findings of an *Xist*-independent effect on growth, we examined our datasets to identify candidate X-genes that are imprinted in the preimplantation embryo. We used a pooled genotype approach to identify genes displaying biased expression from the paternal or maternal X chromosome. Paternal-X biased genes would be expressed consistently across all genotypes excluding X^m^O and X^m^Y, while maternal-X biased genes would be expressed consistently in all genotypes excluding X^p^O and X^p^O *Xist*^Δ^ (Fig. 4a). To ensure that our embryos were developmentally equivalent, we focused our screen on E2.0 and E2.5, prior to when the delay in X^p^O and X^p^O *Xist*^Δ^ embryos initiates. We identified several known imprinted genes, including *Xist*, *Jpx, Ftx*, *Rhox5* and *Fthl17b,e,f* ^40,53,54^, supporting the robustness of our approach (Extended Data Fig. 5a). *Xlr3b* and *Xlr4b/c*, which show X^m^-biased expression in adult tissues^55–57^, did not meet our stringent imprinting criteria in the preimplantation embryo (Extended Data Fig. 5b; see Methods).

**Figure 4.**
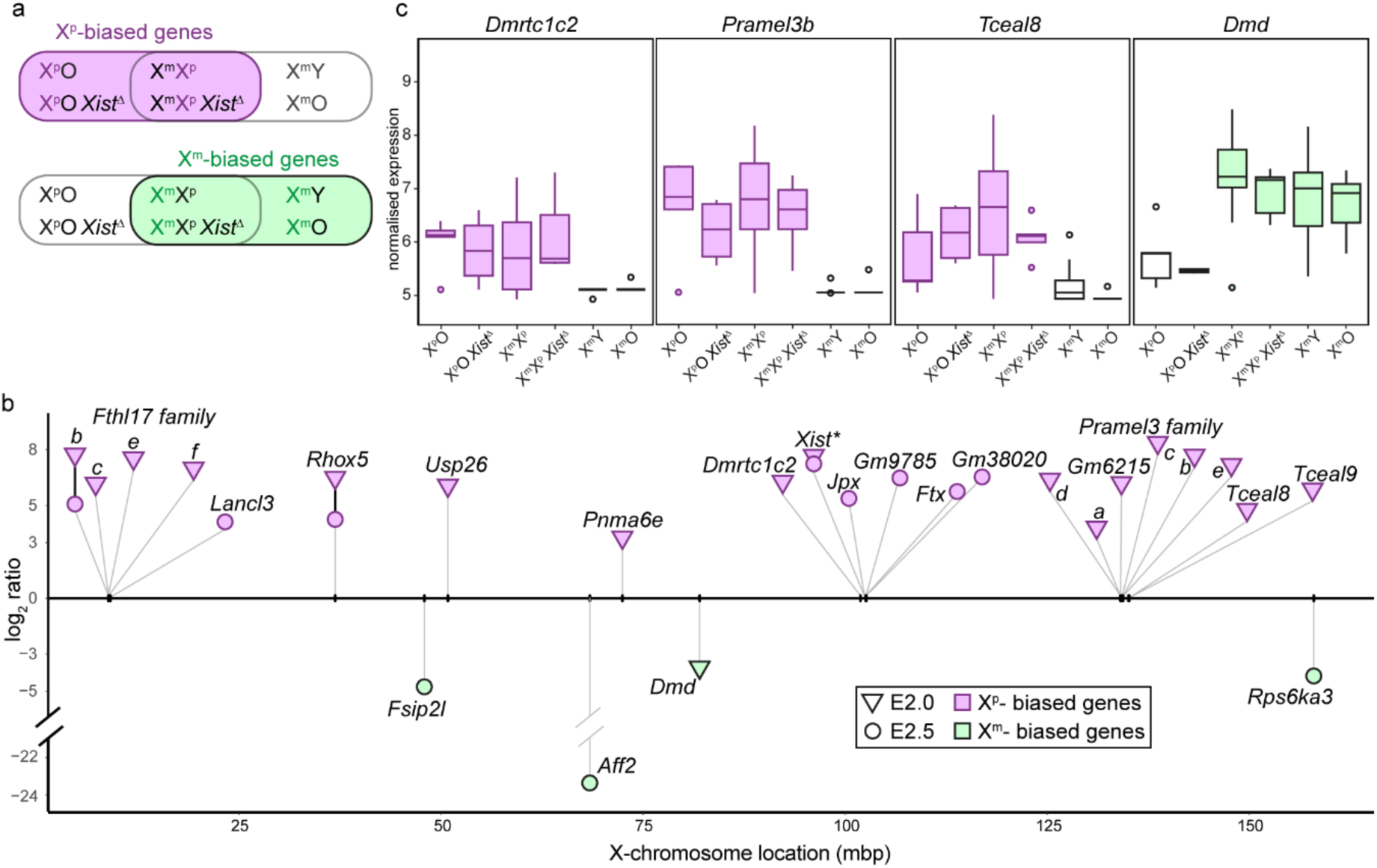
Identification of candidate X-imprinted genes. **a**. Cartoon showing the genotype combinations used to identify X^p^- and X^m^-biased genes in magenta and green, respectively. **b**. X^p^ and X^m^-biased genes shown by position on the X chromosome and log_2_FC at E2.0 (triangle) and E2.5 (circle). Genes showing biased expression at both timepoints are linked with a solid black line. *Paternal-biased expression *Xist* was not observed in our screen. This is expected, because it is deleted in the two *Xist*^Δ^ genotypes, but it is shown for information purposes. **c**. Boxplots of representative X^p^-biased (magenta) and X^m^-biased (green) genes, showing normalised expression across all genotypes at E2.0 (see also Extended Data Fig. 5); *P* values derived from Kolmogorov-Smirnov test.

Using our screen, we identified nineteen novel candidate X-imprinted genes (Fig. 4b, c). These were noteworthy in several regards. First, they showed a bias (15 out of 19 genes) towards being expressed from X^p^. Second, X^p^-expressed genes included several multicopy gene families: beyond the ampliconic *Fthl17* and *Rhox* clusters, we observed X^p^-biased expression of *Pramel3* and *Tceal* cluster genes, as well as the duplicated *Dmrtc1c2* gene (Fig. 4b, c; Extended Data Fig. 5c). The fact that these genes were expressed from X^p^ explained why they were initially identified as female-biased in our X^m^Y and X^m^X^p^ comparison (Fig. 1f). Finally, we noted that many X^p^-biased genes have been reported as expressed predominantly in the male germline, including the *Fthl17*, *Rhox*, *Pramel3*, *Tceal* and *Dmrtc1c2* multicopy genes families, but also single-copy genes *Usp26* ^58,59^ and *Pnma6e* ^60^ (Extended Data Fig. 5c). Our findings reveal thus several potential X-imprinted regulators of early development.

## Discussion

Here we have explored the contribution of sex chromosomes to the preimplantation embryo transcriptome. We find that most sex differences in gene expression are autosomal. Thus, sex chromosomes influence the preimplantation embryo predominantly via *trans* autosomal effects. This influence is even more pronounced in embryos lacking a second sex chromosome, as exemplified in X^m^O, OY and X^p^O embryos. We also find that the trophectoderm transcriptome is heavily influenced by both the Y chromosome and the paternal inactive X chromosome. Our data indicate that this effect is mediated by X-Y homologues in which the X-encoded copy escapes X-inactivation. Strong candidates for this effect include *Utx/y, Kdm5c/d, Ddx3x/y* and *Eif2s3x/y*, which satisfy this criterion and are also deeply conserved regulators of chromatin modification, RNA stability and translation^35,36^.

Our results also reveal the extent of X-imprinting in the early embryo. By removing *Xist*, we identify candidate maternal X-encoded growth promoters and paternal X-encoded growth repressors for future study. We find that, like autosomal imprinted genes^61^, candidate X-imprinted genes are predominantly paternally expressed. The concept of a paternal X-encoded repressor is of interest when considering why imprinted X-inactivation evolved to specifically silence the paternal rather than maternal X chromosome. An imprinted, maternal X-expressed product would be present in both sexes. However, a paternal X-expressed product would be present only in females. If this paternal X-product were a growth repressor, it would selectively disadvantage growth in females. By choosing the paternal X chromosome for silencing, imprinted X-inactivation may prevent this outcome, thereby balancing the growth potential of male and female embryos.

Finally, we show that paternal X-expressed imprinted genes include several multicopy gene clusters that are also expressed in developing male germ cells^41,58^. The significance of this finding warrants future investigation. The X chromosome encodes a disproportionately large number of male germline genes, particularly for those expressed in mitotically-differentiating spermatogonia^58^, and developing spermatids^41^. During male meiosis, the X and Y chromosomes are silenced by meiotic sex chromosome inactivation (MSCI)^62–64^. Subsequently MSCI persists, causing most sex-linked genes to remain repressed during sperm differentiation^41,65,66^. Multicopy and some single-copy X-genes are unusual, exhibiting activation during spermatid differentiation and thus evading the repressive after-effects of MSCI ^41^. The permissive epigenetic state that enables this expression in spermatids may have an enduring effect, leading to ongoing expression after fertilisation when the paternal X is handed down to early female embryos.

---

**Extended Data Figure 1.**
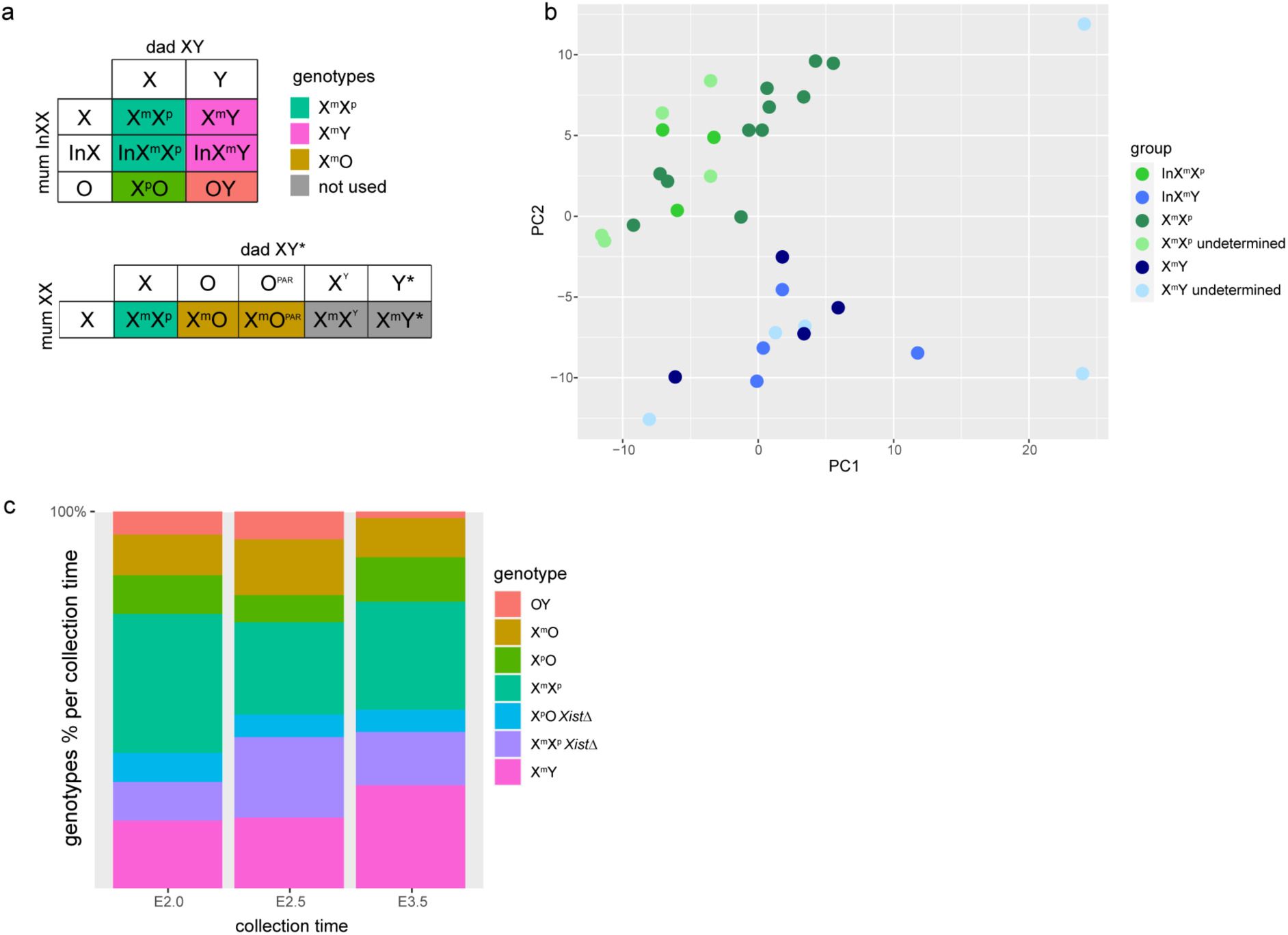
Crosses used in this study. **a.** Table showing crosses used to generate the different genotypes used in this study. The top cross generates two types of X^m^X^p^ embryo (one carrying an inversion on one X chromosome; InX), and two types of X^m^Y embryo, one carrying the same X-inversion. Similarly, the second cross generates two types of XmO embryo, one carrying and one not carrying the tiny pseudoautosomal region. To generate *Xist*^Δ^ mice, the first cross was the same, except the father is X^m^Y *Xist*^Δ^. **b.** Representative PCA of ICM cluster showing that the two X^m^X^p^ genotypes co-cluster relative to X^m^Y genotypes; identification of embryos as X-versus InX-carrying is possible in most but not all embryos (see Materials and Methods). **c.** Bar graph representing the proportion of genotypes by collection time.

**Extended Data Figure 2.**
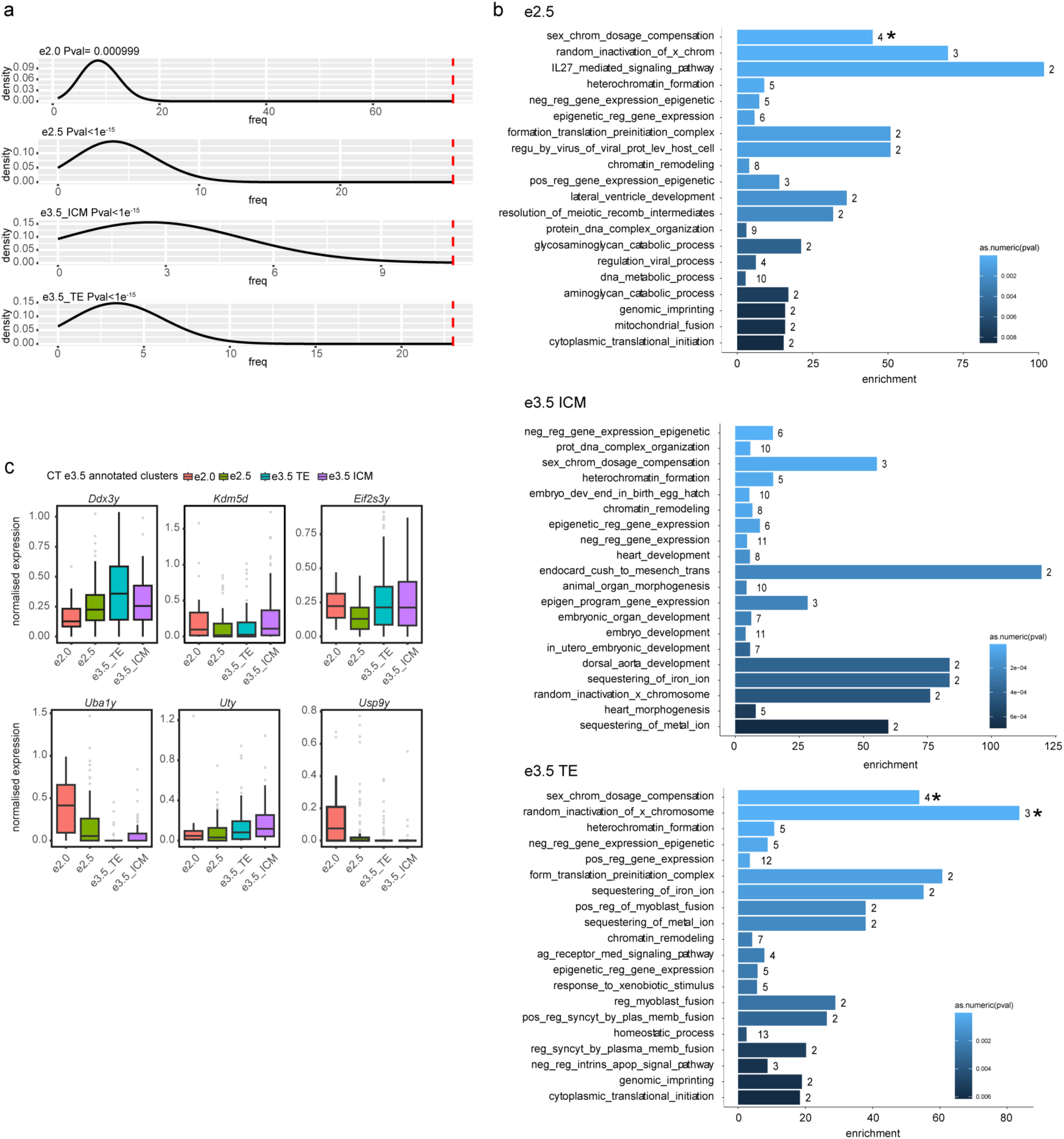
X^m^X^p^ – X^m^Y comparison continued. **a.** Line graph showing over-representation of X-genes in DE analysis. Black line represents expected distribution of DE X-genes and the red dashed line shows the true number of DE X-genes. **b.** Functional enrichment analysis of X^m^X^p^ vs X^m^Y E2.5, E3.5 ICM and TE DE genes; * = p-adjusted p-value <0.05. **c.** Expression of ancestral Y-genes in X^m^Y embryos.

**Extended Data Figure 3.**
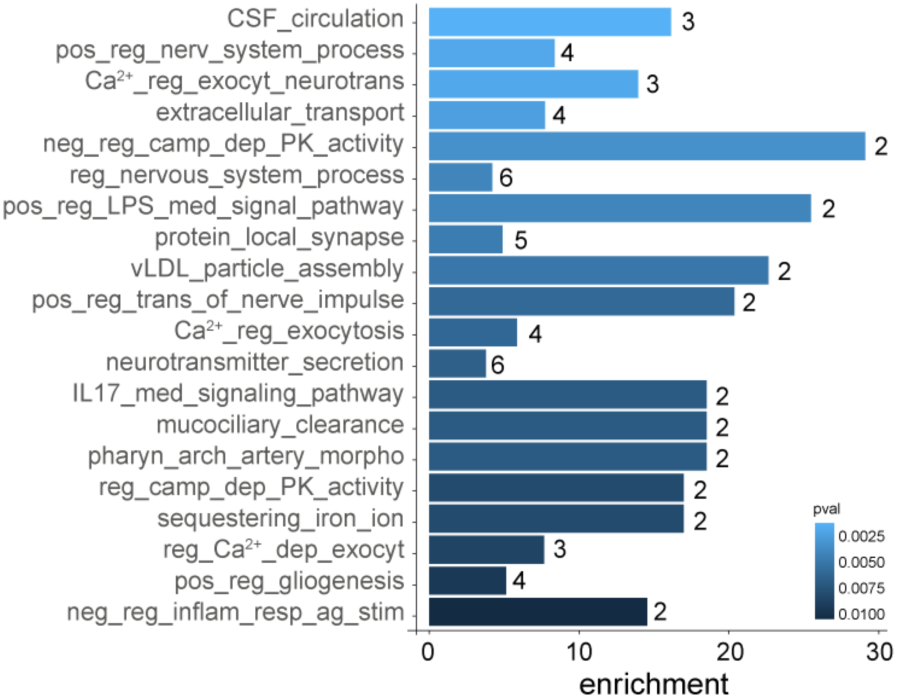
Functional enrichment analysis of overlapping E3.5 TE cluster DE genes between X^m^X^p^ vs X^m^O and X^m^Y vs X^m^O genotypes.

**Extended Data Figure 4.**
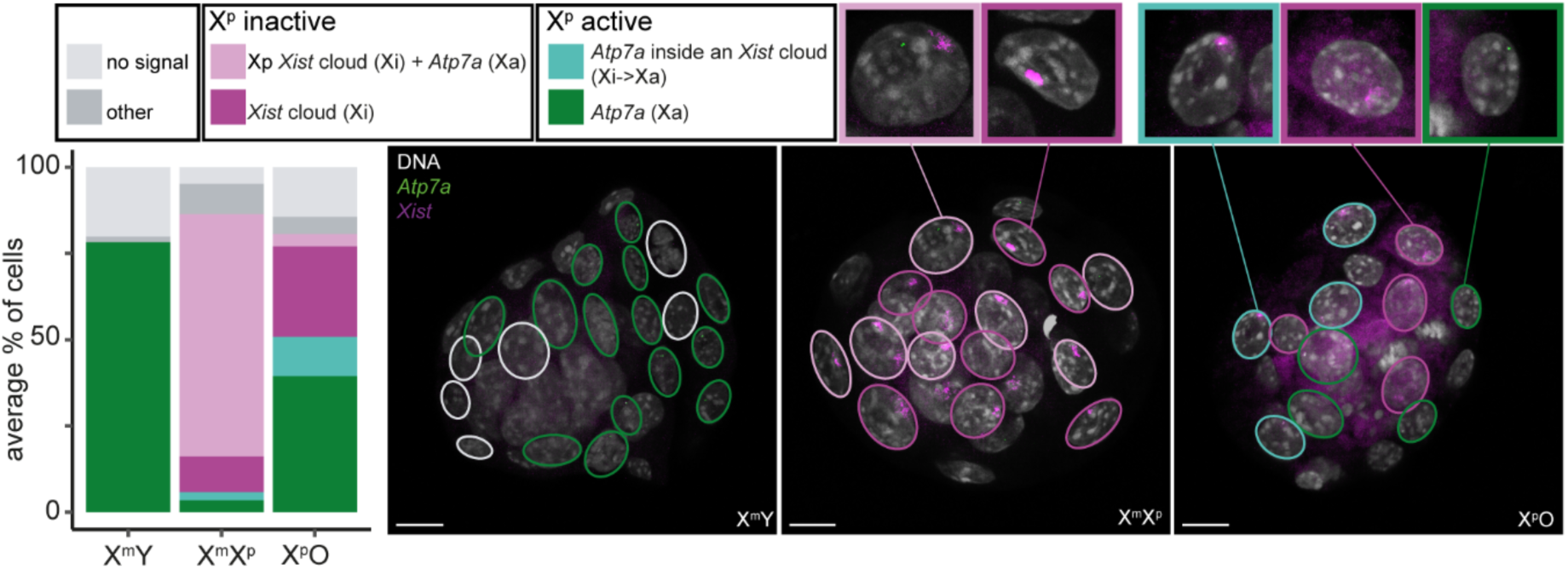
RNA FISH analysis of X-inactivation in X^p^O embryos. Left: bar plot indicating the percentage of cells expressing *Xist* and *Atp7a* in X^m^Y, X^m^X^p^ and X^p^O embryos. Categories: No signal, other (more than one *Xist* cloud or more than 1 *MSN* dot), X^p^ inactive (*Xist* cloud with or without an additional *Atp7a* in another location of the nucleus), X^p^ active (a single *Atp7a* signal, surrounded or not by an *Xist* cloud signal). X^m^Y n= 7, X^m^X^p^ n=7, X^p^O n= 5 embryos. Right: Z-projection of confocal sections (1µm thickness – total thickness of 3.5, 3.5 and 4.5µm respectively) of E3.5 embryos stained for DNA (grey), *Atp7a* (green) and *Xist* (magenta) RNA. Nuclei that are present fully in each projection are encircled, coloured accordingly to their category plotted in the left panel. Scale bar: 20µm.

**Extended Data Figure 5.**
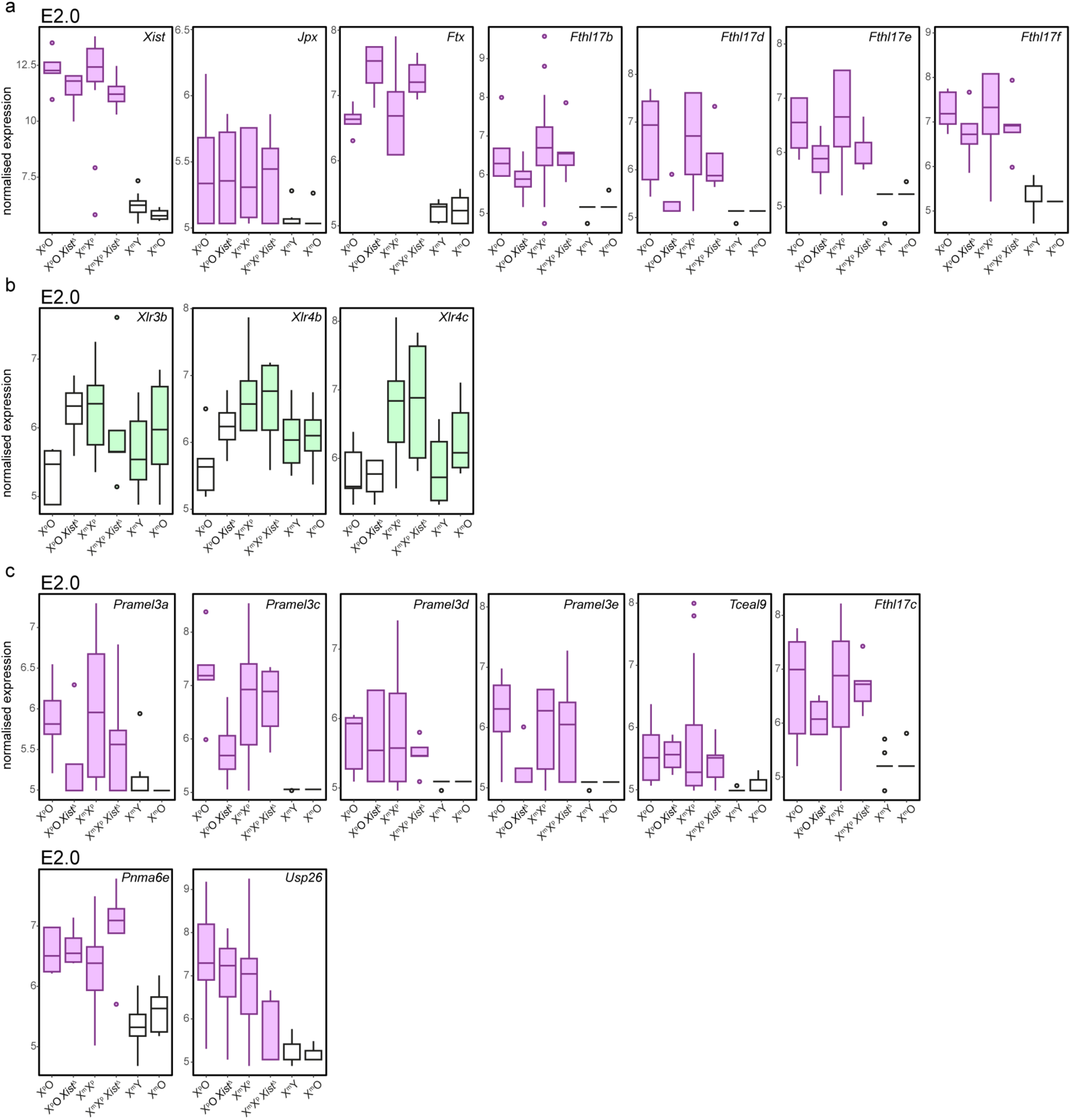
Boxplots of representative X^p^-biased (magenta) and X^m^-biased (green) genes, showing normalised expression across all genotypes at E2.0. **a**. Transcripts described in the literature to be preferentially expressed from X^p^. **b**. Transcripts described in the literature to be preferentially expressed from X^m^. **c**. Transcripts newly described in our study to be preferentially expressed from X^p^.

## Material and methods

### Animals

Mice were maintained in the Francis Crick Institute Biological Research Facility in accordance with United Kingdom Animal Scientific Procedures Act 1986 (Project Licence P8ECF28D9). Mice were housed in individually ventilated cages (GM500, Tecniplast) with a 12:12 light:dark cycle and automatic air and humidity systems. Adults were housed in groups of 3-4 animals, with the sexes housed separately. Mice had free access to water and food and were provided enrichment activities including rodent balls and nesting boxes. Matings for timed collection of embryos were conducted by placing females into the cage of males at approximately 17:00. Observation of a vaginal plug the following morning was taken to indicate mating had occurred between 00:00-02:00. All mice used were generated and maintained on an inbred C57BL/6 background, unless otherwise stated. X^p^O embryos were generated by crossing females heterozygous for an X-linked inversion^29^ with X^m^Y males (Extended Data Fig. 1a, upper). X^m^O embryos were generated by crossing XX females with males carrying a Y-linked rearrangement^30^ (Extended Data Fig. 1a, lower). X^p^O *Xist*^Δ^ embryos were generated by crossing females heterozygous for an X-linked inversion with males carrying an *Xist*^Δ^ allele.

### Embryo and cell collection

For E2.0, embryos were collected at 02:00, ∼48 hours after mating, by flushing oviducts with follicle holding medium (FHM^67^). For E2.5 and E3.5, embryos were collected at 12:00-14:00, ∼60 hours and ∼84 hours after mating, respectively, by flushing uterine horns with FHM. Embryos were then stored at 4°C in PBS for single cell dissociation or fixed directly for immunostaining or RNA FISH. For single-cell dissociation, embryos were incubated in either Acid Tyrode (Merck-Sigma, #T1788) for ∼30s at room temperature or Pronase (5 mg/ml, Millipore, 537088-25KU) for 5 min at 37°C to remove the zona pellucida and then in TripLE (Gibco, 12604013) for 5 min at 37°C to allow dissociation. Embryos were then transferred to a drop of calcium magnesium-free PBS with 5% human serum albumin (SAGE Media, ART-3001) and cells were dissociated by repeated pipetting using glass capillaries. After a wash in PBS, single cells were transferred to individual wells of a 96-well plate containing 2.5 µl of RLT plus buffer (Qiagen, 1053393), frozen on dry ice and stored at −80°C. For E3.5 embryos, duplicates of 2-3 cell pools per embryo were used for genotyping using Illumina SurePlex for whole-genome amplification (see below).

### Post-implantation embryo collection and genotyping

MF1 strain mice were utilised, as described in Thornhill & Burgoyne^9^. Pregnant dams were culled at 12:00 10 days *post coitum*, and the uterine horns removed and placed in ice cold PBS. Individual embryo sacs were dissected free of maternal tissue, and the embryo was freed from the membranes. Each embryo was placed on a 30mm Petri dish and fluid removed with the corner of a lint free tissue, before weighing on a Sartorius Cubis analytical balance MSE225P (Sartorius). Embryos were then frozen in liquid nitrogen before storage at −80°C. DNA was extracted from frozen foetal membranes using PureLink Genomic DNA Mini kit (Thermo Scientific) following the manufacturer’s instructions. The presence of a Y chromosome and In(X) chromosome was determined by PCR^68,69^ (Supplementary Table 2). Chromosome copy number was measured using either qPCR as previously described^70^, or by digital droplet PCR (Bio-rad QX200) with Taqman reagents (Supplementary Table 2), and calculated using QX Manager software.

### G&T-seq

Single cells were prepared for genome and transcriptome sequencing as described previously^71^. Briefly, mRNA and genomic (g) DNA was separated using oligo-dT beads on a Beckman-Coulter liquid-handling platform. RNA-seq libraries were prepared following either a Smart-seq2^72^ or a FLASH-seq low-amplification protocol^73^ utilising Dragonfly and Mosquito HV liquid handlers (SPT Labtech). Sequencing was carried out on a variety of Illumina platforms, including HiSeq4000, NovaSeq 6000 and NovaSeq X+. gDNA underwent whole-genome amplification using SurePlex (Illumina, PR-40-415101-00) following the manufacturer’s instructions at half volume, followed by library preparation by either SQK-LSK109 or SQK-LSK110 and SQK-PCB096 (all Oxford Nanopore Technologies). Libraries were sequenced on a MinION flow cell (either FLO-MIN106D or FLO-MIN114) to a minimum depth of 10000 reads per sample. Basecalling was performed using Guppy, and fastq files were aligned to GRCm38 using nf-core/nanoseq. Using the output of SAMtools idxstats, the proportion of reads was then determined for each sex chromosome from the total number of aligned reads and compared to a reference set comprising data compiled from multiple karyotypically normal female and male samples.

### Immunostaining

Embryos were fixed for 30min in pre-warmed (37°C) fixation buffer at room temperature. Fixation buffer: 1.6% PFA, 100mM HEPES, 50mM EGTA, 10mM MgSO4, 0.2% Tween-20, 0.01% PVP in water. After fixation, they were washed three times in 0.1% Tween20 in PBS for 5 min and permeabilised in 0.2% Tween20 for 30min at room temperature. Embryos were blocked in 0.1% Tween20, 3% BSA in PBS for at least 2 hours at room temperature or overnight at 4°C. They were incubated in the primary antibody overnight at 4°C and incubated in the secondary antibody for two hours at room temperature (shaking). Embryos were imaged in 1% BSA PBS containing 1:2000 Hoechst 33342. Primary antibodies used in this study were: rabbit anti-H2AK119ub1 (Cell Signaling, 8240, 1:500), rabbit anti-Nanog (Cosmo Bio, RCAB001P, 1:500), goat anti-Cdx2 (R&D Systems, AF3665-SP, 1:500). Secondary antibodies against the appropriate species were used as following: anti-rabbit 488 (NANOG) or anti-rabbit 647 (H2AK119ub1), anti-goat Alexa546.

### RNA and DNA FISH

Embryos were fixed in 4% PFA in 0.01% PVP PBS at 4°C for 30min and washed three times for 10min at 4°C in 0.01% PVP PBS. Embryos were permeabilised in −20°C methanol at least overnight, and then hybridised for at least 16 hours at 37°C in hybridisation buffer^74^ containing *Xist* (Fosmid – W112363H9) and Atp7a (BAC – RP23-106F42) probes. Labelling by nick translation was performed with Cy5 and Alexa488 respectively, at a concentration of 1:15. After circular amplification of 0.5µl (10ng) BAC template (Cytiva, 25-6400-50), the BAC was resuspended in 75µL nuclease-free water. 7.5µL of the circular amplified BAC was then used for a 12 hour nick translation reaction (Abbott Molecular) at 15°C with 10µL enzyme, with either ChromaTide Alexa Fluor 488-5-dUTP (Molecular Probes, C11397) or Aminoallyl-dUTP-Cy5 (Jena Bioscience, NU-803-CY5). After RNA FISH, embryos were used for DNA FISH to confirm the number of X and Y chromosomes. Briefly, after washing in 2x SSC PVP and 50% hybridisation buffer/formamide for 5min, embryos were denatured for 10min at 83°C in 50% hybridisation buffer/formamide. Embryos were then incubated for >16 hours in hybridisation solution containing X chromosome paint (XMP X Green – Metasystems, D-1420-050-FI) and Y chromosome paint (XMP Y Orange – Metasystems, D-1421-050-OR) at a concentration of 1:20.

### Imaging and analysis

RNA, DNA FISH and immunostained samples were imaged on a LSM880 confocal microscope (Zeiss), with a 40x water (immunostaining) or 63x oil (RNA and DNA FISH) immersion objective. Images were then analysed using Fiji ^75^ and Imaris (Bitplane) software. Cell number (Fig. 3a) was determined by counting visually all cells in 3D projection in Imaris software, using all different channels: NANOG and CDX2 only are show but embryos were additionally stained for GATA4 and DNA to achieve an exhaustive counting. Whole projection was performed using Fiji software. RNA and DNA FISH (Fig. 3f) were counted using the “cell counter” plugin on Fiji. Plots were generated in R.

### Generating scRNA-seq raw counts & filtering

Raw RNA-seq reads were processed using the RNA-seq nf-core pipeline (v3.14). Genotypes X^m^Y and OY were aligned to the mm39 genome using star_rsem. Genotypes which lacked a Y chromosome (X^m^X^p^, X^p^O, X^p^O *Xist*^Δ^ X^m^X^p^ *Xist*^Δ^ and X^m^O) were aligned to a modified mm39 genome which didn’t include the Y chromosome sequence. The two resulting raw counts tables were merged. Removal of lower quality cells was done using the SingleCellExperiment R package (v1.2.0). Cells were removed if their percentage of mitochondrial reads was greater than four median absolute deviations (MADs). To ensure we retained high quality cells were applied an nFeature cut-off, removing cells which contained nFeatures less than three MADs and removing samples with nCounts in the bottom 5% quantile. Finally, we removed cells with a high percentage of ERCC transcripts (> 3 MADs) that concomitantly showed fewer than 8000 nFeatures.

### Single-cell visualisation

The filtered cells were then processed in the Seurat R package (v4.3.0), using the Seurat v4 scRNA-seq integration workflow. Briefly, CreateSeuratObject() was used to generate a Seurat object, specifying the min.cells=3 parameter. The data was normalised using NormalizeData, followed by FindVariableFeatures. The dataset was scaled using ScaleData. Principle Component Analysis was applied using RunPCA followed by FindNeighbours and FindClusters. Uniform Manifold Approximation and Projection was estimated for the integrated dataset using RunUMAP. Integration was then performed by using SplitObject to split the Seurat object by batch, then running NormalizeData then FindVariableFeatures.

SelectIntegrationFeatures and FindIntegrationAnchors were used to find anchors between the batches, then IntegrateData was used to generate an integrated Seurat object, specifying all genes as features.to.integrate and k.weight=60. The integrated object was then rescaled using ScaleData, followed by RunPCA and RunUMAP. E2.0 and E2.5 samples were treated as their own clusters and annotated as “E2.0” and “E2.5” respectively, for E3.5 we used a 0.2 cluster resolution together with known TE and ICM markers ^76^ to annotate remaining clusters as “E3.5_ICM”, “E3.5_TE” and “E3.5_uncommitted”. Genes used in the X-to-A ratio were identified using a 30% coverage threshold, specifically genes had to be expressed in at least 30% of all samples at >=0.1 expression in the RNA slot. This identified 226 X-linked genes and 7704 autosomal genes. For each cell, the median expression value for the X-linked genes and autosomal genes was calculated using the colMedians function. The X-to-A ratio per cell was the calculated as X-linked median / autosomal_median. Results were visualised as a boxplot.

### Pseudobulk, DESeq2 and functional enrichment

To identify differentially expressed genes between genotypes, a pseudobulk approach was used. Raw counts were pseudobulked using three combined parameters, cluster and genotype and embryo. The pseudobulked counts were generated using the AggregateExpression Seurat function. Each cluster was treated as a separate dataset and was processed in R using the DESeq2 package (v1.40.2). Lowly expressed genes were filtered out by applying a rowSums filter of >=10 to the raw counts table. Principal component analysis (PCA) plots were generated using the top 500 most variable genes, applying the vst() function and limma::removeBatchEffect() to remove batch effects for PCA visualisation. Raw counts were batch corrected and normalised using the DESeq() function, specifying “∼batch + Genotype” in the design formula. Log2 fold change (log2FC) and adjusted p-values between genotypes were calculated using the lfcShrink function, specifying shrinkage method “ashr”. Differentially expressed genes (DEGs) were defined as genes which had an adjusted p-value < 0.05 and log2FC > 0.5. Volcano plots were generated using the EnhancedVolcano package (v1.18). Functional enrichment of differentially expressed genes was done as described ^77^. In order to determine over-representation of X-linked genes (Extended Data Fig. 2a), for each XX vs XY comparison, the number of differentially expressed genes was first randomly sampled from the mouse genome. From the randomly sample gene set, the number of X-linked genes was then calculated. This was repeated 1000 times to generate an expected distribution of X-linked genes per timepoint. The true number of differentially expressed X genes was then compared to the random distribution to determine over-representation

### Imprinting Screen

To identify paternal-X biased genes, raw counts were pseudobulked by into “contains paternal-X” and “does not contain paternal-X” (Fig. 4a, top) as detailed above. The pseudobulk counts were then processed in DESeq2. Potential paternal-X biased genes were identified by using lfcShrink to find X-linked differentially expressed genes that were up-regulated in the “contains paternal-X group”. The same method was done to identify maternal-X biased genes (Fig. 4a, bottom).

To calculate the log2ratio for paternal-X candidates, we took the log2FC for each paternal-X candidate and subtracted from it, the log2FC of same gene from maternal-X comparison. The same methodology was employed to calculate the log2ratio for maternal-X candidates. If the log2FCratio was positive the gene was classified a paternal-X biased, if the log2FCratio was negative it was classified as maternal-X biased.

We then applied a stringent filtering workflow to retain only the high confidence genes. For each paternal-X biased gene, we ensured that all paternal-X genotypes had similar levels of expression. This was done by doing Kolmogorov–Smirnov (KS) test between all combinations of paternal-X genotypes. If the result for each test was insignificant, then the gene passed. Then we repeated the KS test methodology on the remaining two genotypes that did not contain a paternal-X, if the result was also insignificant then the gene passed. Then we filtered the candidates based on the DESeq2 baseMean (average expression) selecting only those that were > 10. Finally, we filtered on the log2FCratio, looking for those genes that had a log2FC ratio > 3. The same filtering workflow was then applied to the maternal-X biased candidates. The paternal and maternal-X biased genes that passed these filtering steps were then visualised in Fig. 4b and 4c.

### Classification of InX

To determine whether a genotype contained an inverted X-chromosome (InX) we used the alignment files generated by nf-core RNA-seq and parsed them through bam-readcount (v0.8.0), specifying 17 X-linked SNPs. Output of bam-readcount was further processed using the bam-readcount accompanying script “brc-parser.py”. SNP metrics were pseudobulked by embryo. Classification of genotypes into InX or X was done using the sum pseudobulked SNV statistic to allow for classification of InX or X. Each locus was called for X or InX if the number of supporting reads was greater than 2; and the embryo was called for X or InX if all classified loci were concordant. For non-concordance, the embryo was “undetermined”.

## Data Availability

scRNA-seq data have been deposited at GEO (GSE281792).

## Acknowledgements

We thank the Francis Crick Institute Biological Research Facility, Genomics Science Technology Platform (STP), Bioinformatics and Biostatistics STP, and High-Performance Computing team, Kathy Niakan, Clare Simon, and members of the J.M.A.T. lab for comments and discussion on the manuscript. Work in the Turner lab is supported by the European Research Council (CoG 647971) and the Francis Crick Institute, which receives its core funding from Cancer Research UK (FC001193), UK Medical Research Council (FC001193) and Wellcome Trust (FC001193). For the purpose of Open Access, the author has applied a CC BY public copyright licence to any Author Accepted Manuscript version arising from this submission.

## Author contributions

D.M.S and J.M.A.T conceived and designed the project. D.S. performed embryo single-cell harvests, G&T-seq, and WGA WGS experiments. W.V performed computational analyses. A.C. performed embryo and single-cell harvests, immunostaining, DNA and RNA FISH, and generated figures. S.M. performed embryo single-cell harvests. R.R. performed image analysis. S.S., P.M. and H.K. performed computational analysis. O.A.O. managed the animal colonies. J.M.A.T wrote the manuscript, and supervised the study with M.N.S.

